# Digital Communication Preferences and Environmental Behavior: A Mixed-Methods Study of Youth Waste Management Education in Buddhist Bhutan

**DOI:** 10.1101/2025.08.10.669562

**Authors:** Smith Boonchutima, Thinley Lhendup, Ibtesam Mazahir

## Abstract

**Background:** Understanding youth perceptions and communication preferences regarding environmental education is crucial for developing effective interventions, particularly in understudied cultural contexts. Most environmental behavior research focuses on Western or large developing countries, leaving significant gaps in understanding small developing nations with unique cultural frameworks.

**Objective:** This study explored youth perceptions, communication preferences, and content effectiveness for waste management education within Bhutan’s Buddhist collectivist culture, addressing identified research gaps in small developing countries.

**Methods:** A convergent mixed-methods design combined quantitative surveys (n=185) with qualitative interviews (n=10) at Yangchenphug Higher Secondary School in Thimphu, Bhutan. The theoretical framework integrated Theory of Planned Behavior, Social Learning Theory, Diffusion of Innovations Theory, and Community-Based Social Marketing principles.

**Results:** Students demonstrated exceptionally high environmental awareness (99.5%) and positive attitudes toward waste separation (mean scores 4.4-4.8 on 5-point scales). Social media emerged as the overwhelmingly preferred communication channel (78.4%), with videos being the most engaging content format (81.6%). Key barriers included time constraints (61.6%), infrastructure limitations (51.9%), and motivational challenges (46.5%). Notably, social influence showed only moderate impact (mean=3.7), lower than typically found in collectivist cultures. Qualitative analysis revealed five themes: shared responsibility burdens, engagement sustainability challenges, system effectiveness limitations, civic values education needs, and institutional support requirements.

**Conclusions:** Bhutanese youth show strong environmental awareness coupled with distinct motivational patterns where environmental behavior appears more individually driven than typically found in collectivist cultures, suggesting Buddhist values of personal responsibility create unique frameworks. Digital communication channels, particularly video content through social media, offer promising avenues for environmental education. A multifaceted approach combining improved infrastructure, educational reinforcement, and culturally-appropriate communication strategies is essential for promoting sustainable waste management practices among youth in similar contexts.

## 1. Introduction

Environmental education and sustainable waste management have become increasingly critical in developing countries experiencing rapid urbanization and changing consumption patterns.

Recent global trends show youth environmental activism rising since 2018, with digital platforms playing increasingly important roles in environmental communication (Pickard et al., 2020). However, significant research gaps exist in environmental behavior studies focusing on small developing countries, where most research has been conducted in Western or larger developing nations (Ifegbesan et al., 2022).

Bhutan, renowned for its commitment to Gross National Happiness (GNH) and environmental preservation, presents a unique case study. The country’s commitment to remaining carbon-neutral and achieving zero waste by 2030 requires active youth participation as future environmental stewards (UNESCAP, 2021). Yet limited research has examined how Bhutan’s distinct Buddhist collectivist culture influences environmental behavior patterns, particularly among youth who represent key agents of environmental change.

### 1.1 Research Gaps in Small Developing Countries

Despite extensive environmental behavior research, significant gaps exist in studies focusing on small developing nations. Most pro-environmental behavior research has been conducted in Western and high-income countries, with limited focus on smaller developing nations where cultural values, economic constraints, and institutional factors may differ substantially (Ifegbesan et al., 2022). This geographic bias leaves critical gaps in understanding unique environmental behaviors and challenges faced by smaller developing countries.

Additionally, insufficient data exists on how socio-demographic variables and psychological dimensions affect pro-environmental behavior in smaller developing countries, as existing studies are often limited to specific regions or larger developing countries (Díaz et al., 2020).

Economic constraints and institutional gaps specifically impact environmental behavior in ways that may be unique to smaller developing nations, where resource limitations and policy implementation challenges create distinct barriers.

### 1.2 Buddhist Cultural Context and Environmental Behavior

Understanding environmental behavior in Bhutan requires consideration of its unique Buddhist collectivist cultural context. Buddhist values significantly influence environmental behavior in South Asian contexts, rooted in core principles such as karma (environmental responsibility), ahimsa (nonviolence), and bodhichitta (compassion for all living beings) (Dorzhigushaeva & Kiplyuks, 2020). Bhutan’s sacred cosmology, influenced by Buddhism and local beliefs, supports environmental conservation and shapes community interactions with the environment (Allison, 2017).

In collectivist cultures, environmental behavior is typically influenced by family and community norms more than in individualistic societies (Morren & Grinstein, 2021). However, Buddhist principles may create unique motivational patterns. Research shows that subjective norms in the Theory of Planned Behavior are particularly strong in Asian contexts, where social harmony and group consensus significantly influence individual decisions (Moon et al., 2021). Yet Buddhist principles such as karma and personal responsibility may create distinct motivational frameworks that differ from typical collectivist societies.

### 1.3 Digital Communication and Youth Environmental Engagement

The digital revolution has fundamentally transformed youth environmental engagement. Social media platforms have proven highly effective in promoting environmental awareness and behavior change, with digital platforms reaching greater numbers and engaging more users compared to traditional media (Hrei et al., 2024). Over 80% of modern digital youth use information and communication technologies to raise environmental awareness (Petrova et al., 2023).

Video content has emerged as particularly effective in environmental education, with research demonstrating youth preferences for video content over traditional materials for environmental learning (Ahmad et al., 2015). Educational videos about environmental topics have shown significant increases in content understanding and improved attitudes toward environmental protection (Kleinhenz & Parker, 2017).

### 1.4 Study Objectives

This study addresses four key research questions within Bhutan’s cultural context: (1) What are the perceptions of Bhutanese youth towards waste separation? (2) What communication channels do Bhutanese youth prefer for receiving waste management information? (3) What types of content are most effective in engaging Bhutanese youth in waste separation practices? (4) How do current waste management practices in Bhutan influence youth participation and engagement?

## 2. Literature Review

### 2.1 Theoretical Framework Integration

Understanding youth environmental behavior requires comprehensive theoretical foundations that account for cognitive, social, and cultural factors. This study integrates four key theoretical frameworks: Theory of Planned Behavior (TPB), Social Learning Theory (SLT), Diffusion of Innovations Theory (DOI), and Community-Based Social Marketing (CBSM).

The integration of multiple behavioral theories in environmental research has shown effectiveness, though implementation faces challenges. Primary motivations for integrating theories include creating more comprehensive explanations of behavior and enhancing behavior change intervention efficacy (Hagger & Hamilton, 2020).

### 2.2 Theory of Planned Behavior in Environmental Contexts

The Theory of Planned Behavior (TPB), developed by Ajzen (1991), remains widely utilized for explaining environmental behavior. According to TPB, behavioral intention is the most significant predictor of behavior, influenced by attitudes toward the behavior, subjective norms, and perceived behavioral control.

Recent meta-analytic studies show TPB consistently predicts environmental behaviors across cultural contexts, though component importance varies (Yuriev et al., 2020). Research reveals significant cultural variations in TPB component predictions. In collectivist cultures, subjective norms tend to have stronger predictive power compared to individualistic cultures, with social norms significantly predicting recycling and waste minimization behaviors (Kaplan Mintz et al., 2019).

Cross-cultural investigations reveal that individuals in countries with better environmental quality tend to have slightly more environmental involvement, but this relationship is mediated by economic circumstances, which have greater independent impact on environmental actions (Freymeyer & Johnson, 2010).

### 2.3 Social Learning Theory and Environmental Education

Social Learning Theory (SLT), proposed by Bandura (1977), posits that people learn behaviors through observation, imitation, and modeling. The theory emphasizes social context importance and peer influence roles in shaping behavior, making it particularly relevant for understanding youth environmental engagement.

Research on motivational strategies for sustaining youth environmental engagement has identified effective approaches. Gamification platforms including social elements such as competition and collaboration can significantly boost engagement and sustained participation (Ouariachi et al., 2020). Teachers play crucial roles in promoting environmental behaviors through SLT principles, with research showing that teachers demonstrating pro-environmental behaviors significantly influence students’ attitudes and behaviors (Öztürk & Pizmony-Levy, 2024).

### 2.4 Community-Based Social Marketing Applications

Community-Based Social Marketing (CBSM), pioneered by McKenzie-Mohr (2011), combines social psychology with marketing techniques to promote sustainable community behavior.

CBSM emphasizes direct engagement, barrier removal, leveraging social norms, providing prompts, and encouraging commitment.

School-based CBSM interventions have shown promising results in promoting sustainable waste practices. Research in Iran demonstrated that CBSM strategies significantly increased recycling behavior among students, with intervention groups showing notable increases compared to control groups (Haghighatjoo et al., 2020).

## 3. Methods

### 3.1 Study Design and Setting

This study employed a convergent mixed-methods approach, combining quantitative surveys with qualitative interviews. The mixed-methods design was chosen based on evidence demonstrating effectiveness in environmental behavior research, particularly for understanding both behavior breadth and cultural meaning depth (Steinmetz-Wood et al., 2019).

Research was conducted at Yangchenphug Higher Secondary School (YHSS) in Thimphu, Bhutan, chosen for its diverse student population representing both urban and rural backgrounds and active environmental initiative engagement. The school’s implementation of programs such as Clean Yangchenphu Captains (CYC) and participation in national initiatives aligned with Clean Bhutan 2030 made it ideal for examining youth environmental behavior within Bhutan’s policy context.

### 3.2 Participants

#### 3.2.1 Quantitative Component

The quantitative sample comprised 185 students aged 18 and above from YHSS. Sample size determination followed established guidelines for environmental behavior studies, where sample sizes between 200-400 are generally considered sufficient (Wolf et al., 2013). Given the exploratory nature of research in this previously unstudied cultural context, this sample size provided adequate power for analyses conducted.

#### 3.2.2 Qualitative Component

The qualitative component involved in-depth interviews with 10 teachers from YHSS, including 3 teachers from upper management, 3 from middle management, and 4 who teach students directly. This purposive sampling strategy ensured diverse perspectives across different organizational levels and teaching experiences.

The data was collected during 15-17 September 2024. Written informed consent was obtained from all participants before their involvement in the study. The questionnaire included a statement informing participants about the purpose of the study, the voluntary nature of their participation, and their right to withdraw at any time without consequences. By proceeding with the questionnaire, participants provided their written consent to take part in the study. To ensure the privacy and confidentiality of the participants, no personally identifiable information was collected during the survey. The data were anonymized and stored securely, with access restricted to the researchers directly involved in the study. The study adhered to the ethical guidelines and principles set forth by Chulalongkorn University and the Faculty of Communication Arts, ensuring the protection of participant privacy and confidentiality throughout the research process. The study was approved by the Research Ethics Review System for Research Involving Human Participants (CU-REC), Chulalongkorn University, Bangkok, Thailand.

### 3.3 Data Collection Instruments

#### 3.3.1 Structured Questionnaire

The questionnaire was designed following best practices for environmental behavior research, incorporating elements from validated instruments including the Recycling Attitude Scale (Ugulu, 2015) and instruments measuring environmental behavior in collectivist cultures. The questionnaire assessed:

- Demographic characteristics
- Knowledge and awareness of waste separation
- Attitudes and perceptions toward waste management
- Social influences and adoption patterns
- Communication channel preferences
- Content preferences
- Barriers and challenges

#### 3.3.2 Semi-Structured Interview Guide

The interview guide covered topics including:

- Current waste management education practices
- Effectiveness of current methods
- Challenges and barriers in teaching waste management
- Student engagement and motivation strategies
- Communication channel preferences
- Recommendations for improvement

### 3.4 Data Analysis

Quantitative data were analyzed using descriptive statistics to summarize demographic characteristics, knowledge levels, attitudes, and preferences. Qualitative data underwent thematic analysis following established procedures, involving initial observations, coding generation, theme development, theme review, and theme definition.

## 4. Results

### 4.1 Participant Characteristics

The study included 185 student participants, with majority being 18 years old (72.4%, n=134). Gender distribution showed 63.8% (n=118) identifying as female, while 36.2% (n=67) identified as male. All participants were Grade 12 students, and 75.7% (n=140) had received formal waste management education prior to the study. (Figure 1)

**Figure 1.**
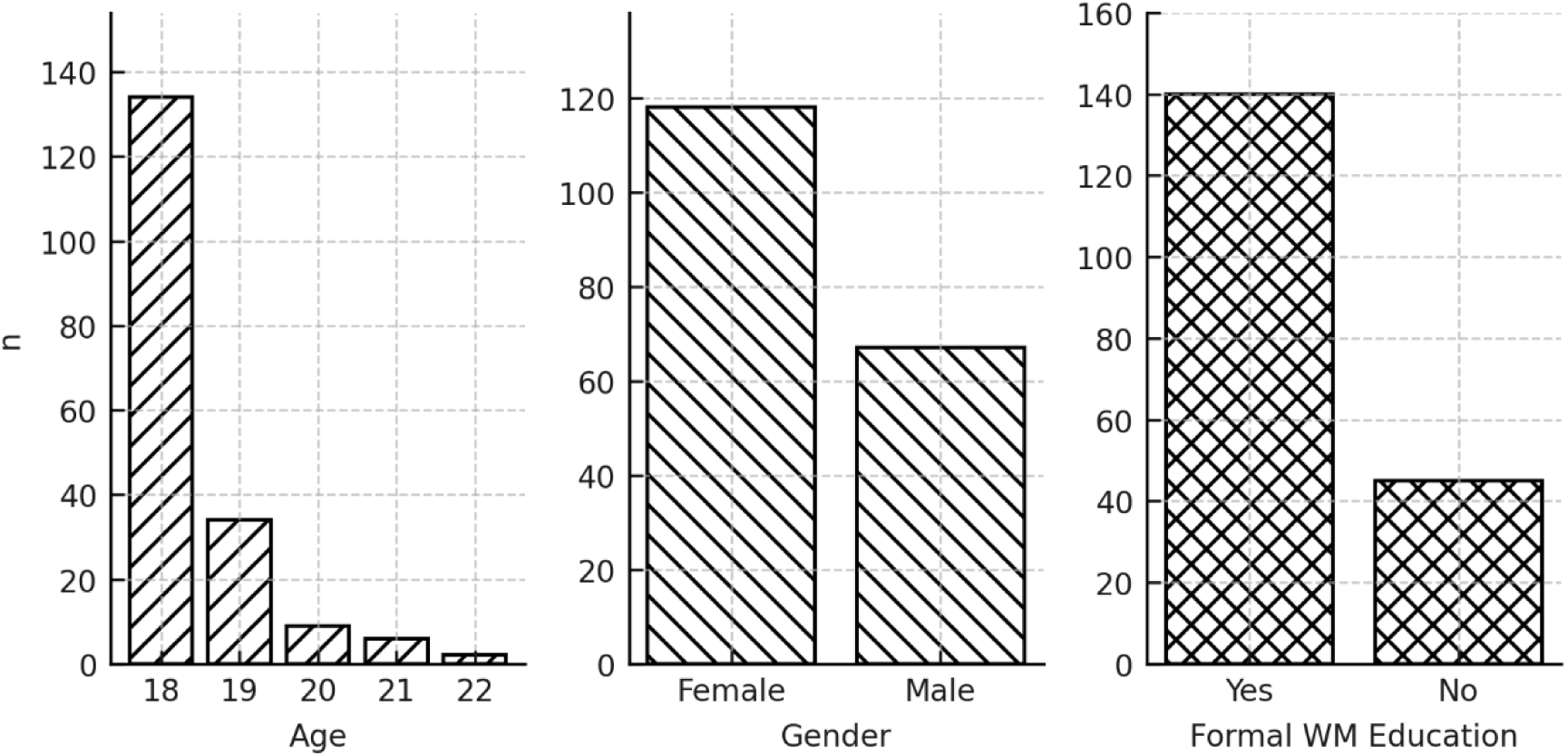
Demographics - Age, Gender, Formal Waste Management Education

### 4.2 Knowledge and Environmental Awareness

Students demonstrated exceptionally high environmental awareness levels, with 99.5% (n=184) reporting awareness of environmental impacts of improper waste disposal. Self-reported waste separation knowledge was moderate (mean=3.0, SD=0.68 on a 4-point scale), suggesting improvement room in technical knowledge despite high general awareness.

Participants demonstrated comprehensive awareness of different waste categories, with 100% awareness reported for all surveyed categories: plastics, paper, glass, organic waste, e-waste, and metals. (Figure 2)

**Figure 2.**
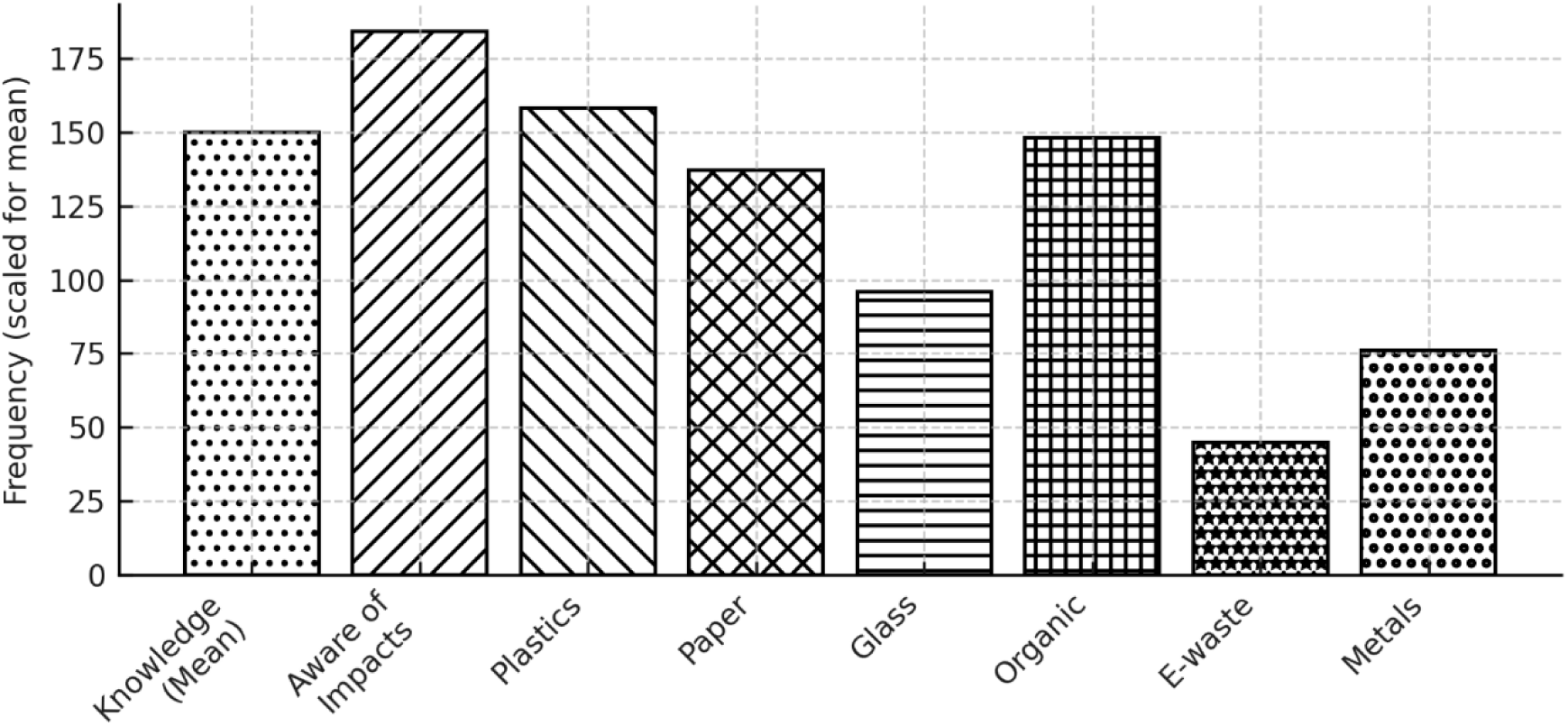
Knowledge levels and waste category awareness

### 4.3 Attitudes and Perceptions

Students showed overwhelmingly positive attitudes toward waste separation:

- **Environmental Importance**: Strong agreement that waste separation protects the environment (mean=4.8, SD=0.52)
- **Personal Responsibility**: High sense of personal responsibility for waste separation (mean=4.5, SD=0.61)
- **Individual Efficacy**: Strong belief that individual efforts make a difference (mean=4.4, SD=0.66)
- **Social Influence**: Moderate influence from friends and family (mean=3.7, SD=0.87)
- **Confidence**: High confidence in waste separation ability (mean=4.1, SD=0.69)

Interestingly, social influence from friends and family showed only moderate impact (mean=3.7), which in collectivist cultures typically shows stronger effects. This finding challenges assumptions about collectivist motivation patterns and suggests environmental behavior in Bhutan reflects Buddhist individualistic responsibility synthesis with collectivist social awareness. (Figure 3)

**Figure 3.**
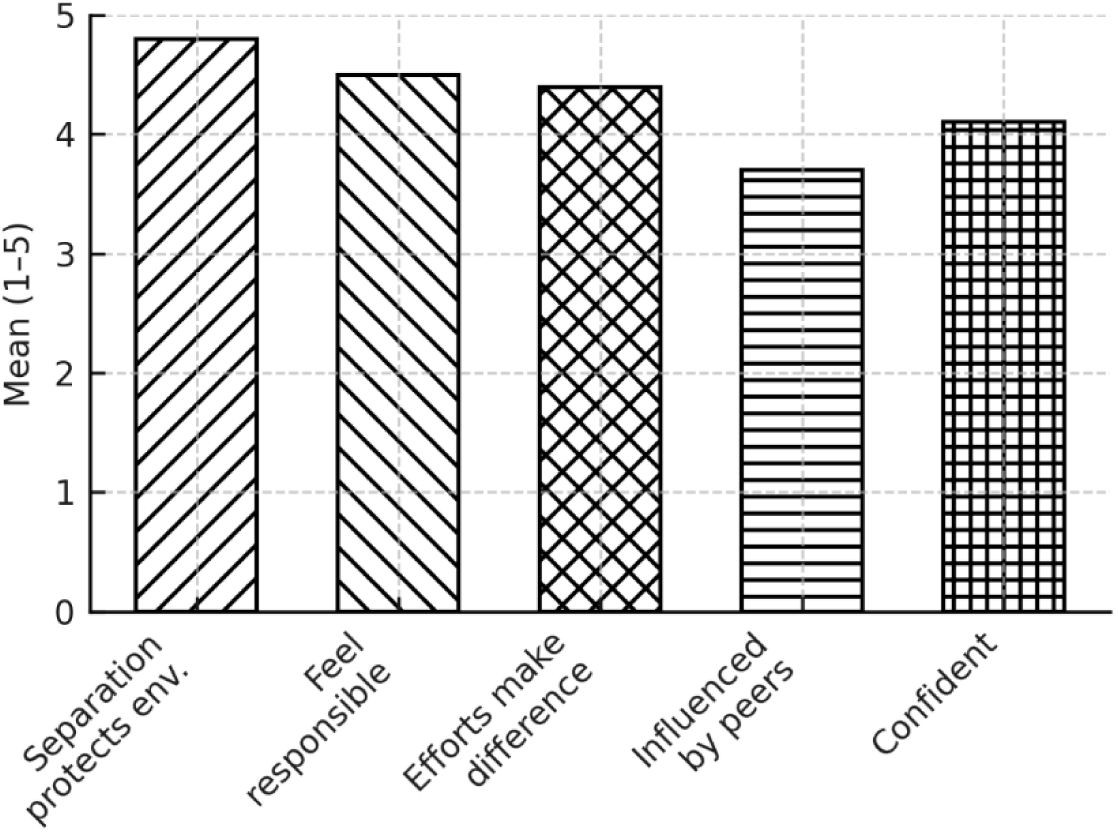
Mean scores for attitudes - environmental importance, personal responsibility, etc.

### 4.4 Communication Channel Preferences

Communication channel preferences revealed strong alignment with global youth digital engagement trends:

#### Preferred Communication Channels

- Social media: 78.4% (n=145) - overwhelmingly preferred
- School announcements: 59.5% (n=110)
- Community events: 41.6% (n=77)
- Traditional media (TV, Radio): 38.4% (n=71)
- Environmental clubs: 31.4% (n=58) (Figure 4)

#### Preferred Content Types

- Videos: 81.6% (n=151) - most engaging format
- Workshops: 43.2% (n=80)
- Posters: 35.7% (n=66)
- Interactive apps: 24.3% (n=45)
- Infographics: 10.8% (n=20) (Figure 5)

Students rated social media campaigns as highly effective for promoting waste separation among youth (mean=4.0, SD=0.82), and 88.1% (n=163) expressed willingness to participate in school or community-based campaigns. (Figure 6)

**Figure 4.**
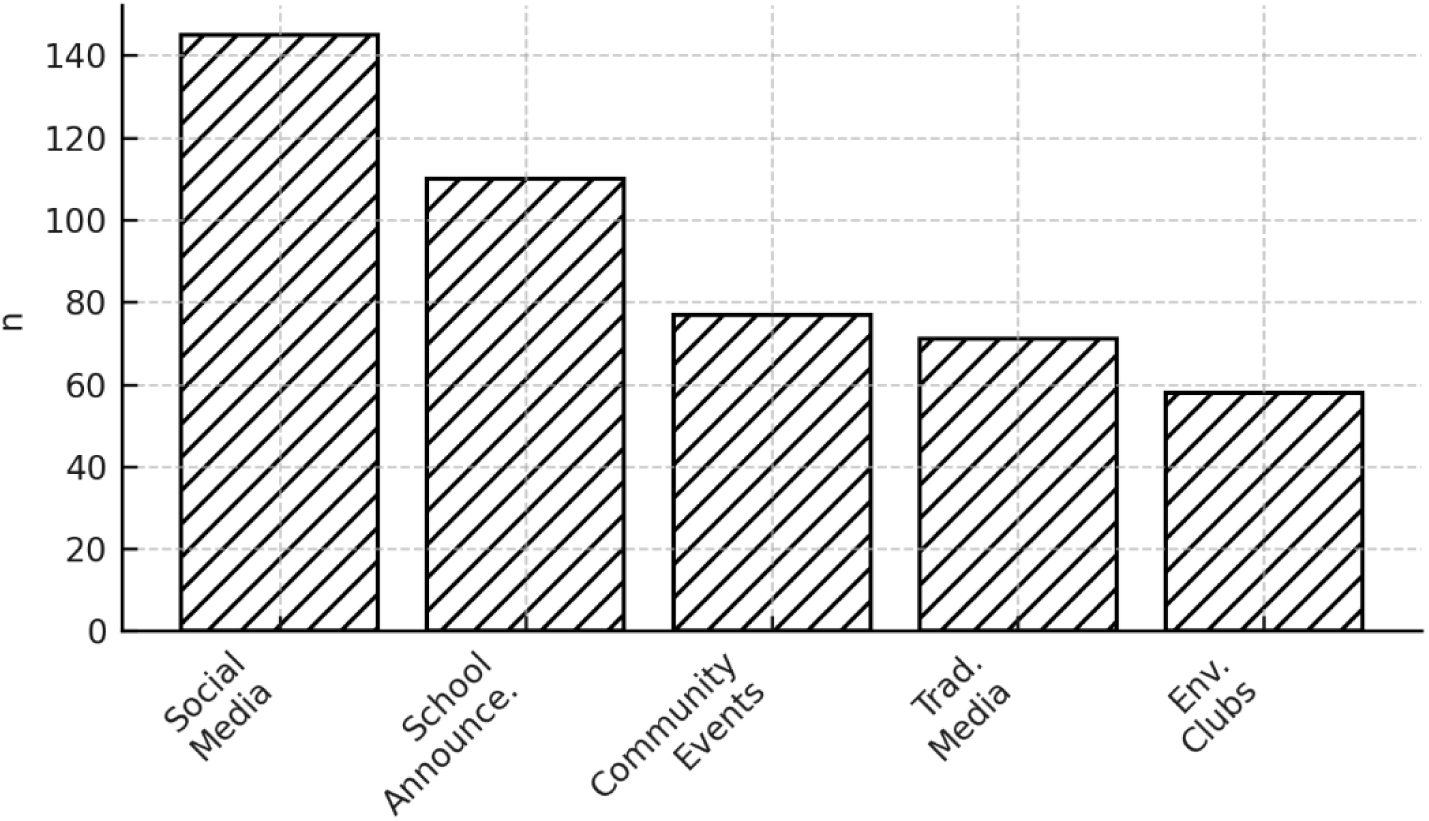
Preferred communication channels

**Figure 5.**
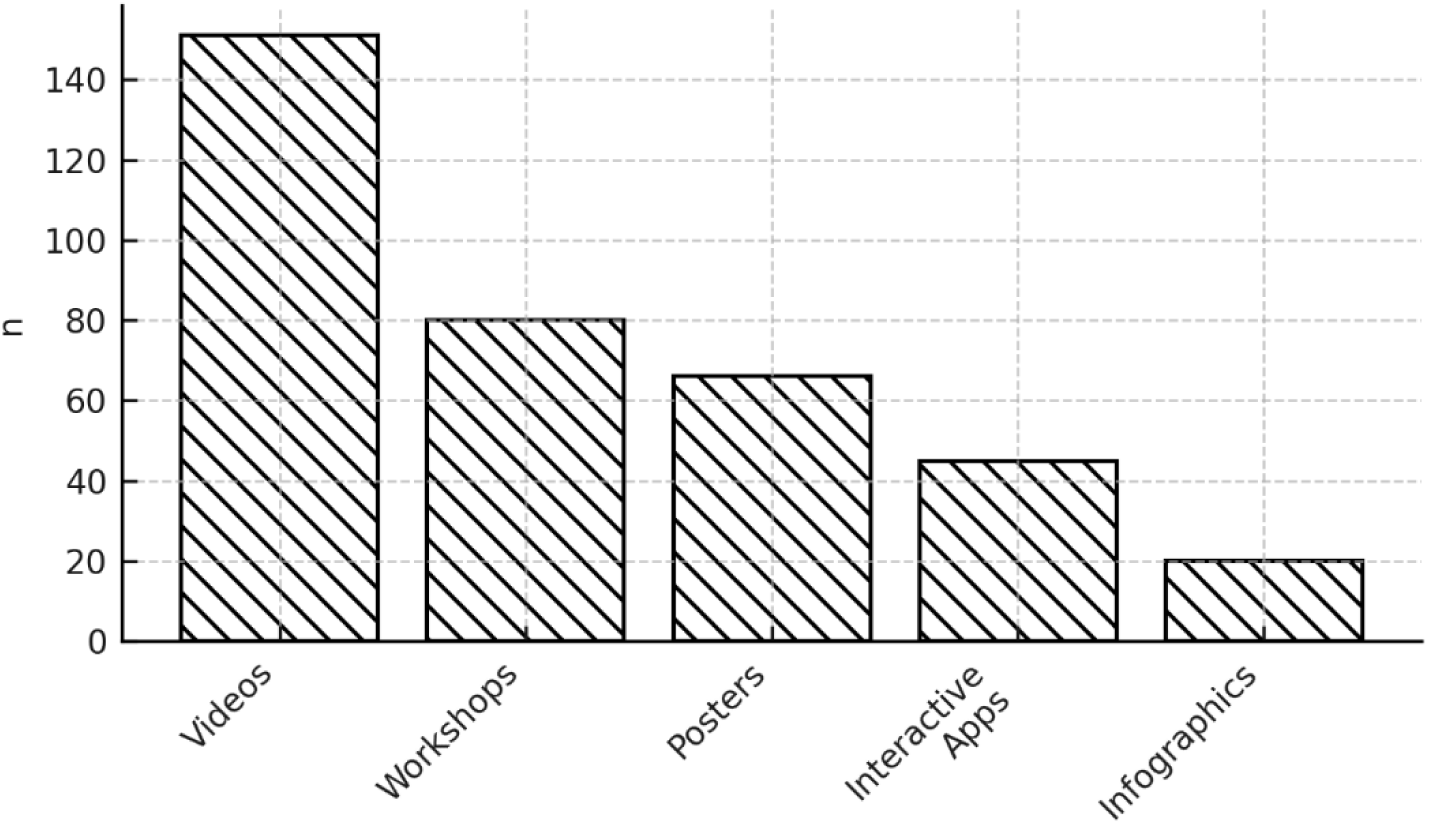
Preferred content types

**Figure 6.**
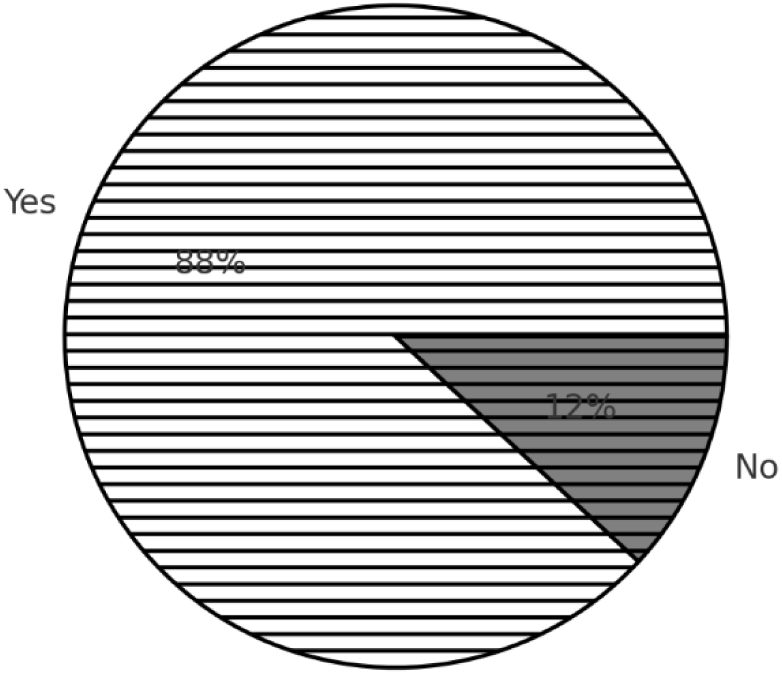
Willingness to participate in campaigns

### 4.5 Adoption Patterns and Innovation Factors

The study revealed important patterns regarding students’ adoption of new waste management practices and factors that encourage their participation in environmental initiatives.

#### Adoption Characteristics

Students demonstrated moderate levels of early adoption compared to their peers (mean=3.5, SD=0.75 on a 5-point scale), suggesting they adopt new practices at a similar pace to their peer group rather than being early innovators. However, willingness to try new waste separation practices was notably high (mean=4.0, SD=0.62), indicating strong openness to environmental innovations despite moderate adoption speed. (Figure 7)

**Figure 7.**
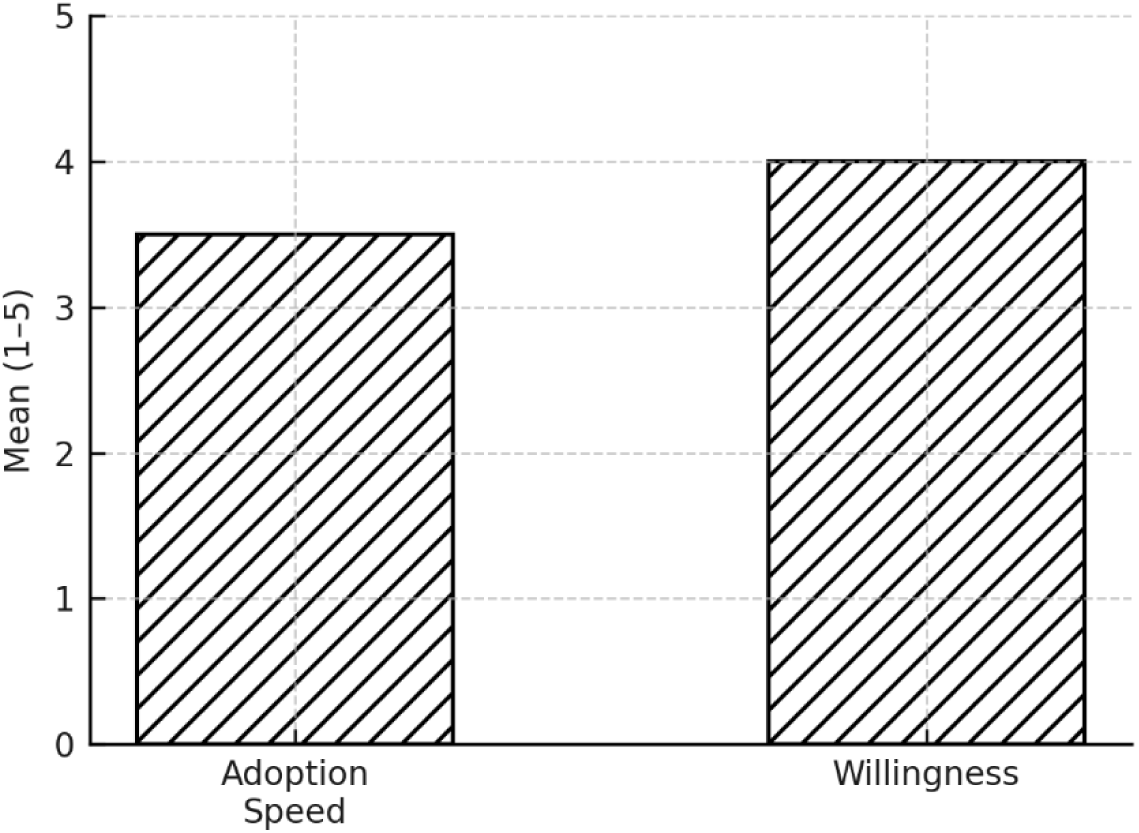
Mean scores for Adoption speed and willingness to try new waste separation practices

Regarding actual participation, 45.4% (n=84) of respondents reported having participated in new waste management programs introduced at their school, while 54.6% (n=101) had not yet engaged with such initiatives. This moderate participation rate suggests room for improvement in program reach and engagement strategies. (Figure 8)

**Figure 8.**
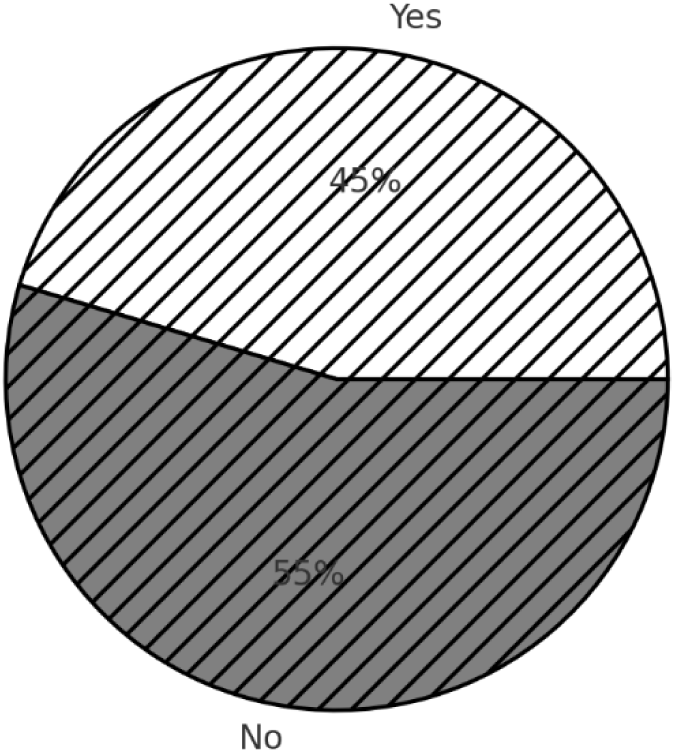
Experience in participating in new waste management programs

#### Factors Encouraging Adoption

When asked about factors that would encourage adoption of new waste separation practices, students identified several key motivators (multiple selections allowed):

- Clear benefits: 67.03% (n=124) - most frequently cited factor
- Seeing others participate: 47.57% (n=88)
- Visible positive outcomes: 41.08% (n=76)
- Ease of use: 25.95% (n=48)
- Trial opportunities: 16.22% (n=30)

These findings align with Diffusion of Innovations Theory, where perceived relative advantage (clear benefits) and observability are key factors in adoption decisions. The importance of “seeing others do it” validates Social Learning Theory’s relevance, emphasizing the role of peer modeling in environmental behavior adoption.

### 4.5 Barriers and Motivational Factors

#### Primary Barriers

- Time constraints: 61.6% (n=114) - most significant barrier
- Lack of facilities: 51.9% (n=96)
- Lack of motivation: 46.5% (n=86)
- Inconvenience: 38.9% (n=72)
- Lack of knowledge: 29.7% (n=55) (Figure 9)

#### Primary Motivators

- Environmental impact: 81.1% (n=150) - strongest motivator
- Personal responsibility: 76.2% (n=141)
- Social approval: 65.4% (n=121)
- Educational incentives: 56.8% (n=105)
- Ease of practice: 54.1% (n=100) (Figure 10)

### 4.6 Student-Suggested Solutions for Barrier Removal

Students provided comprehensive suggestions for overcoming identified barriers to waste management participation. Analysis of open-ended responses revealed multiple solution categories, demonstrating students’ sophisticated understanding of both challenges and potential interventions (Figure 9).

**Figure 9.**
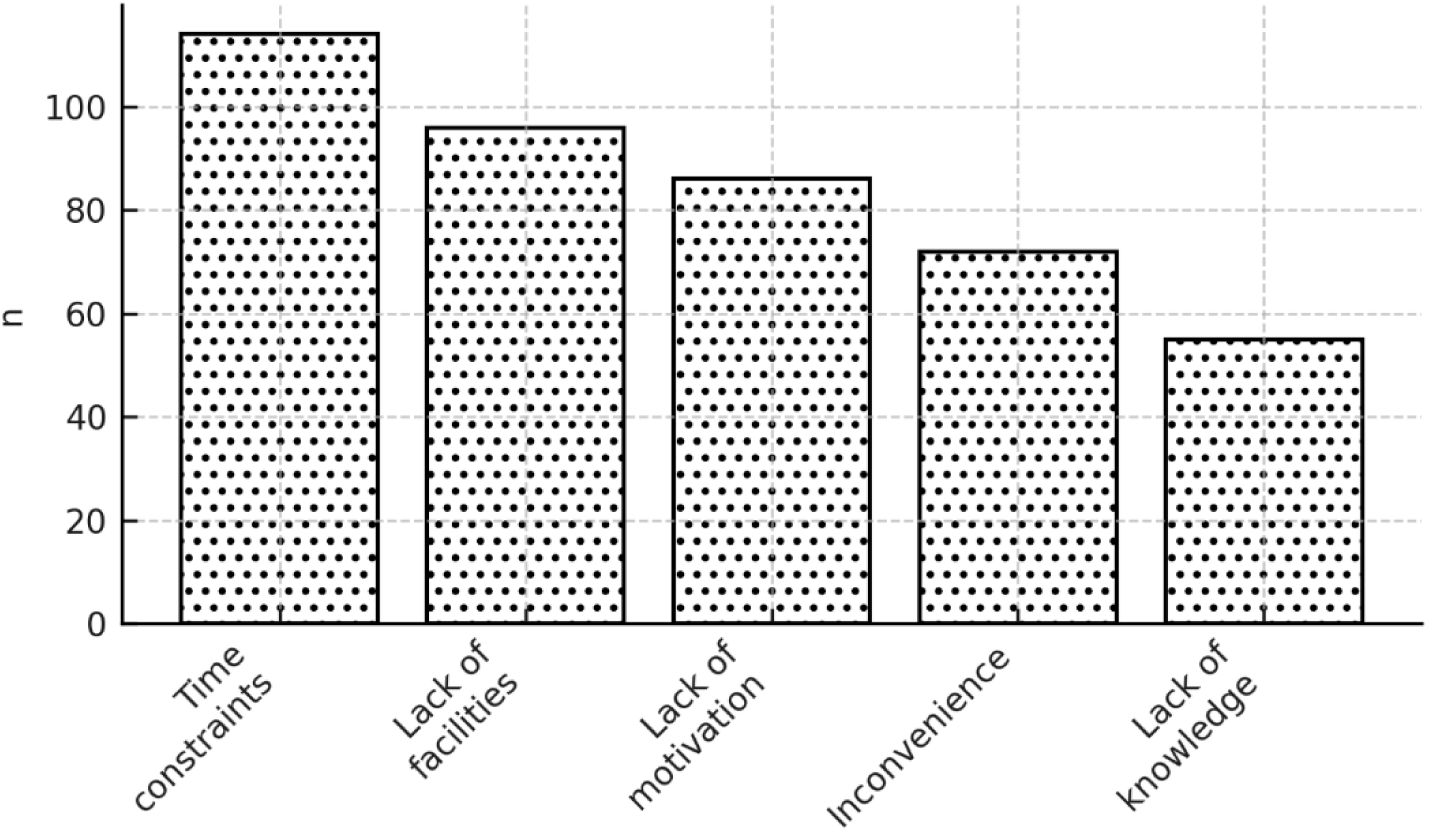
Barriers to Participation in Waste Management

**Figure 10.**
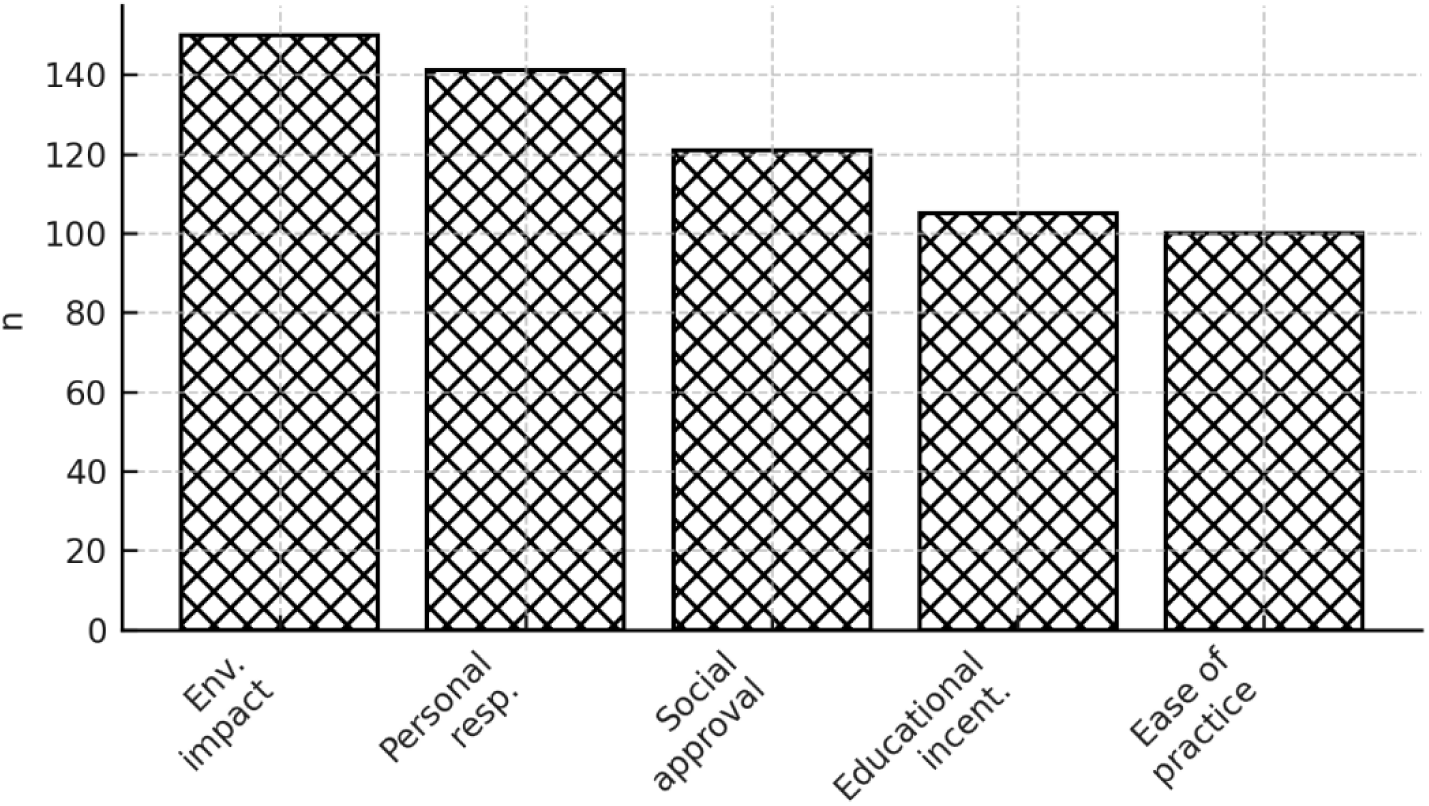
Motivational factors

The most frequently mentioned solution category was education and awareness (55 mentions), with students emphasizing the need for enhanced educational programs and awareness campaigns to promote proper waste separation practices. This was followed by calls for improved infrastructure and facilities (42 mentions), highlighting the critical need for accessible waste management resources and equipment.

These student-generated solutions align with Community-Based Social Marketing principles, emphasizing barrier removal and community engagement as key strategies for promoting sustainable behaviors. The comprehensive nature of these suggestions indicates that youth possess valuable insights into effective intervention design and should be included in program development processes. (Figure 11)

**Figure 11.**
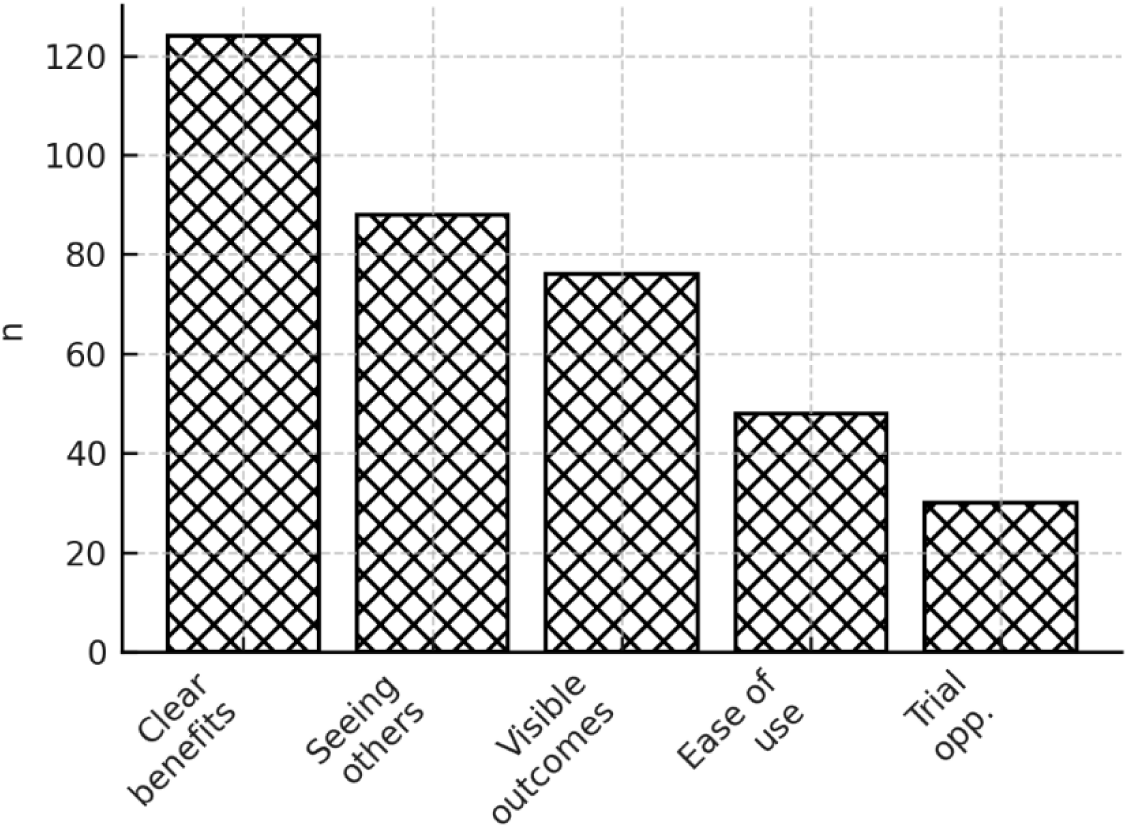
Solution categories by frequency

### 4.6 Qualitative Findings: Teacher Perspectives

Thematic analysis revealed five key themes:

#### 4.6.1 Shared Responsibility and Teacher Burden

Teachers expressed concerns about managing both academic duties and waste management responsibilities. Despite initiatives like Clean Yangchenphu Captains (CYC), much workload still falls on teachers. One participant noted: “Being a class teacher is already a major role… sometimes we might not be giving much attention to waste management, but I do ensure that the class is clean.”

#### 4.6.2 Challenges in Sustaining Student Engagement

While external motivators like prizes temporarily engage students, fostering intrinsic motivation proves challenging. A teacher observed: “It’s a little burdensome for [the CYC captains] because the rest of the classmates leave it up to them.”

#### 4.6.3 Effectiveness of Waste Management Systems

Current systems including waste segregation and competitions show mixed results. A subject teacher noted: “Despite the efforts from the teachers, there still seems to be a lot of gaps between what we tell students and what they do.”

#### 4.6.4 Instilling Civic Values through Education

Teachers emphasized waste management should be viewed as core civic responsibility rather than extracurricular activity. A senior teacher said: “We need to instill in our children the importance of taking care of their own waste… they need to understand the implications it will have on the environment.”

#### 4.6.5 Policy and Institutional Support Needs

While the school has made improvements through initiatives aligned with Clean Bhutan 2030, teachers expressed need for stronger institutional backing. A senior administrator mentioned: “The school has improved a lot since we introduced class-based segregation, but we still face challenges because individuals fail to understand the importance of proper waste management.”

## 5. Discussion

### 5.1 Key Findings Integration

This mixed-methods study reveals that Bhutanese youth demonstrate exceptionally high environmental awareness (99.5%) and positive attitudes toward waste separation, coupled with strong willingness to engage when appropriate support is provided. However, significant practical barriers persist, including time constraints (61.6%), infrastructure limitations (51.9%), and motivational challenges (46.5%).

The study addresses identified research gaps in environmental behavior studies in small developing countries, where most research has focused on Western or larger developing nations (Ifegbesan et al., 2022). The findings demonstrate that environmental behavior patterns in small developing countries like Bhutan may differ from those in larger nations due to unique cultural, institutional, and economic factors.

### 5.2 Cultural Context Analysis: Buddhist Values and Environmental Behavior

The high personal responsibility scores (mean=4.5) likely reflect Buddhist principles of karma and individual accountability for actions affecting others and the environment. The emphasis on environmental impact as the strongest motivator (81.1%) aligns with Buddhist concepts of interdependence and compassion for all beings.

However, the relatively moderate social influence (mean=3.7) challenges typical collectivist culture patterns where social norms usually show stronger effects. This finding suggests that environmental behavior in Bhutan may be more individually motivated than predicted by cultural dimensions theory, indicating that Buddhist values of personal responsibility create distinct motivational frameworks for environmental behavior.

Research on Buddhist environmental ethics shows that these principles create unique frameworks for environmental stewardship that differ from Western environmental ethics (Choudhury, 2025). The synthesis of Buddhist individualistic responsibility with collectivist social awareness creates a unique cultural pattern worthy of further investigation.

### 5.3 Digital Communication Implications

The overwhelming preference for social media (78.4%) and video content (81.6%) aligns with global research on youth digital engagement while providing specific guidance for environmental communication in Bhutan. These findings support research demonstrating social media’s effectiveness in promoting environmental awareness and behavior change among youth (Hrei et al., 2024).

The effectiveness of video content aligns with research showing videos significantly increase student content understanding and improve attitudes toward environmental protection (Kleinhenz & Parker, 2017). However, it’s important to note that while digital communication can effectively influence environmental intentions, translating these intentions into actual behavior changes remains challenging (Huang et al., 2025).

### 5.4 Theoretical Framework Validation

#### 5.4.1 Theory of Planned Behavior

The findings strongly support TPB’s applicability in Bhutan’s cultural context, with high scores on attitudes (mean=4.8) and perceived behavioral control (mean=4.1). However, the moderate influence of social norms (mean=3.7) is lower than typically found in collectivist cultures, suggesting environmental behavior in Bhutan may be more individually motivated than predicted.

#### 5.4.2 Social Learning Theory

The importance of “seeing others do it” as an adoption factor (47.57%) and challenges with peer responsibility in the CYC system validate SLT’s relevance. The qualitative finding that students rely on CYC captains rather than modeling their behavior suggests peer influence requires more systematic implementation.

#### 5.4.3 Community-Based Social Marketing

The identification of specific barriers (time, facilities, motivation) and preferences for direct engagement through workshops strongly supports CBSM’s barrier-focused approach. The high willingness to participate in campaigns (88.1%) combined with practical barriers suggests CBSM interventions addressing infrastructure and time management could be highly effective.

### 5.5 Practical Implications and Recommendations

#### 5.5.1 Educational Strategy Development

**Digital-First Approach**: The study provides strong evidence for prioritizing social media platforms and video content in environmental education initiatives. Environmental education has essential effects on students’ waste sorting behavior, and videos are more effective than traditional materials in promoting environmental awareness (Jorge et al., 2024).

**Theory-Based Interventions**: The positive attitudes and strong sense of personal responsibility suggest that educational interventions based on TPB could be highly effective, as demonstrated in recent studies showing significant improvements in waste separation behaviors among primary school students (Fallah-Nejad et al., 2023).

#### 5.5.2 Infrastructure and Policy Recommendations

**Facility Improvements**: The high frequency of facility-related barriers (51.9%) indicates that environmental behavior change requires supportive infrastructure. Research demonstrates that convenience of waste sorting facilities significantly impacts behavior (Hao et al., 2020).

**Time Management Integration**: Rather than treating environmental activities as additional tasks, integration into daily academic routines could address the primary barrier of time constraints (61.6%). This approach aligns with research showing successful school-based environmental programs that incorporate environmental education into existing frameworks.

### 5.6 Limitations

The study focused on a single school in Thimphu, which may not represent all Bhutanese youth, particularly those in rural areas with different infrastructure and cultural contexts. Cultural specificity means findings may be specific to Bhutan’s unique Buddhist collectivist culture and may not apply to other developing countries with different cultural contexts.

Survey responses may reflect social desirability bias, particularly on environmental attitudes and behaviors in a culture with strong environmental values. The study measured stated preferences and intentions rather than actual behavior, which research shows can differ significantly.

### 5.7 Future Research Directions

Future research should examine long-term effectiveness of interventions based on these findings, particularly whether digital communication strategies and barrier removal approaches lead to sustained behavior change over time. Comparative studies examining environmental behavior across different Buddhist cultures could identify universal versus culture-specific factors.

Research should also address gaps in digital communication effectiveness for environmental education, investigating the intention-behavior gap and developing strategies to bridge this divide in digital environmental education.

## 6. Conclusions

This comprehensive mixed-methods study provides valuable insights into youth perceptions and preferences regarding waste management education in Bhutan, contributing to both environmental behavior theory and practical environmental education applications. The research reveals a promising foundation of high environmental awareness and positive attitudes among Bhutanese youth, while identifying specific barriers and preferences that can guide intervention development.

### 6.1 Theoretical Contributions

The study validates the applicability of multiple behavioral theories (TPB, SLT, DOI, CBSM) within Bhutan’s Buddhist collectivist cultural context while revealing unique patterns that challenge assumptions about collectivist motivation. The finding that environmental behavior appears more individually motivated than typically found in collectivist cultures suggests Buddhist values of personal responsibility may create distinct motivational patterns.

This research contributes to filling identified gaps in environmental behavior studies in small developing countries, demonstrating that environmental behavior patterns may differ significantly from those in larger nations due to unique cultural, institutional, and economic factors.

### 6.2 Practical Applications

The research provides specific, actionable guidance for environmental education:

**Communication Strategy**: Social media platforms using video content should be prioritized, supported by institutional channels and community engagement. The high effectiveness ratings provide clear direction for resource allocation in environmental communication.

**Educational Interventions**: Theory-based educational programs, particularly those using TPB, should be implemented following successful models demonstrated in other contexts. The Health Promoting Schools model could also be effective, given its demonstrated success in improving recycling behaviors.

**Barrier Removal**: Infrastructure improvements and time management integration are essential for translating positive attitudes into consistent behaviors. The specific barriers identified provide concrete targets for intervention development.

### 6.3 Global Relevance

While culturally specific to Bhutan, this research provides insights relevant to environmental education in other developing countries with strong environmental commitments and collectivist cultural values. The systematic approach to understanding youth perceptions, communication preferences, and content effectiveness provides a replicable framework for environmental education research in similar contexts.

The finding that positive attitudes and willingness to engage can coexist with practical barriers that limit participation has implications for environmental education worldwide. The emphasis on infrastructure and institutional support alongside attitude formation challenges education-only approaches to environmental behavior change.

### 6.4 Final Recommendations

Based on comprehensive findings, we recommend:

1. **Immediate Actions**: Develop social media campaigns using video content that emphasize environmental impact and personal responsibility while addressing practical barriers through infrastructure improvements.
2. **Medium-term Development**: Implement systematic peer education programs that expand beyond current CYC systems to engage broader student participation in environmental initiatives.
3. **Long-term Integration**: Work toward systematic integration of environmental education into formal curricula while building institutional capacity and support for sustained environmental education efforts.
4. **Research Continuation**: Conduct longitudinal studies to evaluate intervention effectiveness and comparative studies to understand cultural factors affecting environmental behavior across different contexts.

The research demonstrates that Bhutanese youth possess the awareness, attitudes, and willingness necessary for effective environmental stewardship. By addressing practical barriers and leveraging communication preferences identified in this study, environmental educators can more effectively engage young people in sustainable waste management practices, contributing to broader environmental conservation goals and providing a model for youth environmental engagement in similar cultural contexts globally.

## Ethical Approval Statement

To ensure the privacy and confidentiality of the participants, no personally identifiable information was collected during the survey. The data were anonymized and stored securely, with access restricted to the researchers directly involved in the study. The study adhered to the ethical guidelines and principles set forth by Chulalongkorn University and the Faculty of Communication Arts, ensuring the protection of participant privacy and confidentiality throughout the research process. The study was approved by the Research Ethics Review System for Research Involving Human Participants (CU-REC), Chulalongkorn University, Bangkok, Thailand.

## Acknowledgments

Since, this research is derived from a Master’s thesis, the authors would like to express their sincere appreciation to the thesis committee—Assistant Professor Dr. Papaporn Chaihanchanchai, Ph.D. (Chairperson) and Dr. Ser Shaw Hong, D.B.A. (Examiner)—for their valuable feedback and constructive guidance throughout the research process. They also extend their gratitude to Associate Professor Dr. Preeda Akarachantachote, Ph.D., Dean of the Faculty of Communication Arts, for their continued support and encouragement.

Special thanks are due to the faculty and staff of Yangchenphu Higher Secondary School (YHSS) for their cooperation and assistance during data collection. The authors are also grateful to Khun Sally and Dr. Mao for their personal support, and to the ASEAN & Non-ASEAN Scholarship Program for providing the financial assistance that made this research possible.

## Data availability statement

The data that support the findings of this study are available on request from the corresponding author.

